# High genetic loading of schizophrenia predicts poor response to lithium in patients with bipolar disorder: A polygenic score and cross-trait genetic analysis

**DOI:** 10.1101/209270

**Authors:** Azmeraw T. Amare, Klaus Oliver Schubert, Liping Hou, Scott R. Clark, Sergi Papiol, Urs Heilbronner, Franziska Degenhardt, Fasil Tekola-Ayele, Yi-Hsiang Hsu, Tatyana Shekhtman, Mazda Adli, Nirmala Akula, Kazufumi Akiyama, Raffaella Ardau, Bárbara Arias, Jean-Michel Aubry, Lena Backlund, Abesh Kumar Bhattacharjee, Frank Bellivier, Antonio Benabarre, Susanne Bengesser, Joanna M. Biernacka, Armin Birner, Clara Brichant-Petitjean, Pablo Cervantes, Hsi-Chung y, Caterina Chillotti, Sven Cichon, Cristiana Cruceanu, Piotr M. Czerski, Nina Dalkner, Alexandre Dayer, Maria Del Zompo, J. Raymond DePaulo, Bruno Étain, Peter Falkai, Andreas J. Forstner, Louise Frisen, Mark. A Frye, Janice M. Fullerton, Sébastien Gard, Julie. S Garnham, Fernando. S Goes, Maria Grigoroiu-Serbanescu, Paul Grof, Ryota Hashimoto, Joanna Hauser, Stefan Herms, Per Hoffmann, Andrea Hofmann, Stephane Jamain, Esther Jiménez, Jean-Pierre Kahn, Layla Kassem, Po-Hsiu Kuo, Tadafumi Kato, John Kelsoe, Sarah Kittel-Schneider, Sebastian Kliwicki, Barbara König, Ichiro Kusumi, Gonzalo Laje, Mikael Landén, Catharina Lavebratt, Marion Leboyer, Susan. G Leckband, Alfonso Tortorella, Mirko Manchia, Lina Martinsson, Michael J. McCarthy, Susan McElroy, Francesc Colom, Marina Mitjans, Francis. M Mondimore, Palmiero Monteleone, Caroline M. Nievergelt, Markus M. Nöthen, Tomas Novák, Claire O’Donovan, Norio Ozaki, Urban Ösby, Andrea Pfennig, James B. Potash, Andreas Reif, Eva Reininghaus, Guy. A Rouleau, Janusz. K Rybakowski, Martin Schalling, Peter R. Schofield, Barbara. W Schweizer, Giovanni Severino, Paul. D Shilling, Katzutaka Shimoda, Christian Simhandl, Claire. M Slaney, Alessio Squassina, Thomas Stamm, Pavla Stopkova, Mario Maj, Gustavo Turecki, Eduard Vieta, Julia Volkert, Stephanie. H Witt, Adam Wright, Peter. P Zandi, Philip. B Mitchell, Michael Bauer, Martin Alda, Marcella Rietschel, Francis. J McMahon, Thomas G. Schulze, Bernhard T. Baune

## Abstract

**Importance:** Lithium is a first-line mood stabilizer for the maintenance treatment of Bipolar Disorder (BPD). However, the efficacy of lithium varies widely, with a non-response rate of up to 30%. Biological response markers and predictors are lacking.

**Objective:** Genetic factors are thought to mediate lithium treatment response, and the previously reported genetic overlap between BPD and schizophrenia (SCZ) led us to test whether a polygenic score (PGS) for SCZ could predict lithium treatment response in BPD. Further, we explored the potential molecular underpinnings of this association.

**Design:** Weighted SCZ PGSs were computed at ten p-value thresholds (P_T_) using summary statistics from a genome-wide association study (GWAS) of 36,989 SCZ cases, and genotype data for BPD patients from the Consortium on Lithium Genetics (ConLi^+^Gen). For functional exploration, we performed a cross-trait meta-GWAS and pathway analysis, combining GWAS summary statistics on SCZ and lithium treatment response.

**Setting:** International multicenter GWAS.

**Participants:** Patients with BPD who had undergone lithium treatment were genotyped and retrospectively assessed for long-term treatment response (n=2,586).

**Main outcome measures:** Clinical treatment response to lithium was defined on both the categorical and continuous scales using the ALDA score. The effect measures include odds ratios (ORs) and the proportion of variance explained (R^2^), and a significant association was determined at p<0.05.

**Results:** The PGS for SCZ was inversely associated with lithium treatment response in the categorical outcome (p=8×10^−5^), at P_T_ <5×10^−2^. Patients with BPD who had low polygenic load for SCZ responded better to lithium, with ORs for lithium response ranging from 3.46 [95%CI: 1.42-8.41 at 1^st^ decile] to 2.03 [95%CI: 0.86-4.81 at the 9th decile], compared to the patients in the 10^th^ decile of SCZ risk. In the cross-trait meta-GWAS, 15 genetic loci that may have overlapping effects on lithium treatment response and susceptibility to SCZ were identified. Functional pathway and network analysis of these loci point to the HLA complex and inflammatory cytokines (TNFα, IL-4, IFNγ) as molecular contributors to lithium treatment response in BPD.

**Conclusions and Relevance:** The study provides, for the first-time, evidence for a negative association between high genetic loading for SCZ and poor response to lithium in patients with BPD. These results suggest the potential for translational research aimed at personalized prescribing of lithium.

**Key Points:** *Question:* Does a polygenic score for Schizophrenia (SCZ) predict response to lithium in patients with Bipolar Disorder (BPD)? What are the molecular drivers of the association between SCZ and lithium treatment response?

*Findings:* We found an inverse association between genetic loading for SCZ risk variants and response to lithium in patients with BPD. Genetic variants in the HLA region on chromosome 6, the antigen presentation pathway and markers of inflammation (TNFα, IL-4, IFNγ) point to molecular underpinnings of lithium treatment response in BPD.

*Meaning:* In patients with BPD, an assessment of a polygenic load for SCZ risk variants may assist in conjunction with clinical data to predict whether they would respond to lithium treatment.

## INTRODUCTION

Bipolar Disorder (BPD) is a severe and often disabling psychiatric condition, characterized by recurrent dysregulation of mood with episodes of mania and depression. With an early disease onset and an estimated lifetime prevalence of 1%^1^ to 4.4%^2^, BPD is associated with high personal impairment and societal costs, accounting for 9.9 million years of life lived with disability worldwide^3^, and substantially increased all-cause mortality and risk of suicide^4^. The etiology of BPD is complex, and both genetic and environmental factors have been shown to contribute to the pathogenesis of the disorder^5^. The estimated heritability of BPD ranges from 60% to 85%^6^, and candidate gene^7^ and genome-wide association studies (GWASs)^8–12^ have successfully identified genetic loci implicated in the illness. However, only a small fraction of the heritability is accounted for by replicated genetic variants that have been identified so far^7^.

Lithium stabilizing properties were discovered by Australian psychiatrist John Cade back in 1949^13^. Since then, it has retained a status as the ‘gold standard’ mood stabilizer^14,15^, possessing unique protective effects against both manic and depressive episodes^16^, as well as for suicide prevention^17^. Consequently, lithium is recommended as first-line maintenance treatment for BPD by several clinical practice guidelines^18–21^. However, there is significant inter-individual variation between lithium treatment responders and non-responders. About 30% of patients are only partially responsive, and more than a quarter show no clinical response at all^22^. While clinical studies report a combination of demographic and clinical characteristics as potential predictors of treatment response in patients with BPD^23^, genetic factors also appear to be highly involved^22,24–26^. So far, three GWASs have successfully identified single nucleotide polymorphisms (SNPs) associated with lithium treatment response in BPD pointing to different genetic loci ^22, 27, 28^. To improve the understanding of the molecular mechanisms underlying the therapeutic effects of lithium, alternative genomic approaches can complement GWAS deserve consideration. One such approach is polygenic analysis, which quantifies the combined effects of genetic variants across the whole genome on a given clinical outcome, computed as a weighted summation of effect sizes of multiple independent polymorphisms. An accurate and successful polygenic model may assist early screening for disease risk, clinical diagnosis, and the prediction of treatment response and prognosis. In the current study, we aimed to investigate whether BPD patients with high trait genetic susceptibility for schizophrenia (SCZ), expressed by their SCZ polygenic score (PGS), would respond better or more poorly to lithium compared to BPD patients with a low PGS for SCZ. Additionally, we set out to explore the genetic and molecular underpinnings of any identified association between SCZ and lithium treatment response. A number of previous observations motivated this approach. First, there is increasing evidence for a substantial genetic overlap between BPD and SCZ. The Psychiatric Genomics Consortium (PGC) estimated a shared genetic variation of ~68%, which is the highest among all pairs of psychiatric diagnoses^27^. Consistent with this, several shared risk genes and shared biological pathways associated with both disorders have been identified^28,29,30^, and current sample sizes for SCZ far exceed those available for BPD and thus are better powered. Second, despite these genetic and molecular commonalities, lithium is not an effective medication for people suffering from SCZ^31^, and increased SCZ trait loading in those with BPD might be expected to serve as a predictor for poor treatment response. An earlier family study found an association between family history of schizophrenia and poor response to lithium^32^ Third, during acute illness episodes, BPD and SCZ are often difficult to distinguish clinically because of overlapping psychotic symptoms such as hallucinations, delusions, and disorganization, as well as some common behavioral disturbances such as irritability or anger ^33^. Aiming to predict response to lithium, which could potentially confer advantages for patients and their treating physicians^34^ we sought to evaluate the aggregated effect of genome-wide SNPs for SCZ on lithium treatment response in BPD using a polygenic score approach that was based on the results of the largest SCZ GWAS to date^35^. Further, in order to explore potential genetic and molecular drivers of any detected association, we carried out a cross-trait GWAS meta-analysis, combining the summary statistics from the largest available GWAS for both SCZ^35^ and lithium response^22^. Overlapping SNPs that met genome-wide significance in the meta-GWAS were subsequently analyzed for biological context using the Ingenuity^®^ Pathway Analysis platform (IPA^®^).

## METHODS AND MATERIALS

### Study Samples

#### The International Consortium on Lithium Genetics (ConLi^+^Gen)

The ConLi^+^Gen Consortium (www.ConLiGen.org) is an initiative by the National Institute of Mental Health (NIMH) and the International Group for the Study of Lithium-Treated Patients (IGSLI) (www.IGSLI.org) that was established with the aim of discovering genetic variants responsible for lithium treatment response in BPD^36^. The ConLi^+^Gen study involved patients with BPD from Europe, South America, USA, Asia, and Australia^22^ who had been treated with lithium at some stage since diagnosis. The first GWAS based on this initiative was published in 2016^22^. For the current study, genetic and clinical data collected from 2,586 patients with BPD who were part of the ConLi^+^Gen consortium were analyzed^22,36^. A series of quality control procedures were implemented on the genotype data before and after imputation as described below.

#### Genotyping and quality control

The genome-wide genotypes, as well as clinical and demographic data, were collected by 22 participating sites. Quality control (QC) procedures were implemented using PLINK^37^.

Samples with low genotype rates <95%, sex inconsistencies (X-chromosome heterozygosity), and genetically related individuals were excluded. We also excluded SNPs that had a poor genotyping rate (<95%), an ambiguity (A/T and C/G SNPs), a low minor allele frequency (MAF<1%), or that showed deviation from Hardy-Weinberg Equilibrium (p<10^−6^).

#### Imputation

The genotype data passing QC were imputed on the Michigan server^38^ (https://imputationserver.sph.umich.edu) separately for each genotype platform using the 1000 Genomes Project Phase 3 (Version 5) reference panel. During the imputation process, we used the European reference panel for all the samples except for those from Japan and Taiwan, for which the East Asian reference population was used. After excluding the low-frequency SNPs (MAF<10%); low-quality variants (imputation INFO < 0.9); and indels, the imputed dosages were converted to best guess genotypes. The subsequent polygenic analyses were performed using the best guess genotypes.

#### Discovery GWAS summary data

The PGSs were calculated using the approach previously described by the International Schizophrenia Consortium^39^. This method requires discovery and target datasets. The discovery data, which refers to the GWAS summary statistics-effect sizes (beta, a log of odds ratio), were obtained from a previously published SCZ GWAS^35^ that was publicly available for download by the Psychiatric Genomics Consortium (PGC) http://www.med.unc.edu/pgc/, accessed on March 18, 2017.

#### Target outcome

Lithium treatment response in BPD was defined for patients who had received lithium for a minimum of 6 months. Lithium treatment outcome was assessed using the “Retrospective Criteria of Long-Term Treatment Response in Research Subjects with Bipolar Disorder” scale, also known as the ALDA scale ^40,41^. The ALDA scale is a well-validated tool to rate symptom improvements after treatment with lithium in BPD, and it has shown excellent inter-rater reliability^42^. The ALDA scale quantifies symptom improvement over the course of treatment (A score, range 0–10), which is then weighted against five criteria (B score) that assess confounding factors, each scored 0, 1, or 2. The total score is calculated by subtracting the total B score from the A score, and negative scores are set to zero^22^. We developed two main outcomes for lithium response (categorical and continuous outcome). The categorical (i.e., good versus poor) response to lithium in BPD was defined based on the total score as a cut-off score of 7, in which patients with a total score of 7 or higher were categorized as “responders”. The ALDA score on subscale A was used as a continuous outcome after excluding individuals with a total B score greater than 4 or who had missing data on the total scores of ALDA subscale A or B^22^. In addition to the ALDA scale scores, information on covariates such as age and gender was collected, and further details can be found in an earlier publication^22^.

#### Polygenic scoring

Quality-controlled SNPs were clumped for linkage disequilibrium based on GWAS association p-value informed clumping using r^2^ = 0.1 within a 250-kb window to create a SNP-set in linkage equilibrium using PLINK software run on Linux (*plink --clump-p1 1 -- clump-p2 1 --clump-r2 0.1 --clump-kb 250*). Then, the SNPs at ten p-value thresholds (<1×10^−4^, <1×10^−3^, <0.01, <0.05, <0.1, <0.2, <0.3, <0.4, <0.5, <1) were selected to compute the SCZ PGSs in the ConLi^+^Gen sample. The major histocompatibility complex region was excluded from the PGS calculation because of its complex linkage disequilibrium structure. A genome-wide weighted SCZ PGS for each participant was calculated at each p-value threshold (P_T_) as the sum of independent SNPs genotype dosage (from 0 to 2) of the reference allele in the ConLi^+^Gen genotype data, multiplied by SCZ GWAS effect sizes for the reference allele, estimated as log (OR) divided by the total number of SNPs in each threshold.

### STATISTICAL ANALYSES

For statistical analyses, we applied PGS association analyses, cross-trait meta-GWAS, and Ingenuity Pathway Analysis (IPA) of the cross-trait findings. The details for each analysis are described below.

#### Polygenic score association analysis

Once the PGSs were constructed, the association of the PGSs at each P_T_ and lithium treatment response was evaluated using regression models. While a binary logistic regression was implemented for the categorical outcome (response versus non-response), a linear regression was applied to lithium treatment response on the continuous scale. Using the PGS at the most significant threshold (P_T_ <5×10^−2^), we divided the study samples into ten deciles (1^st^ to 10^th^), ranging from the lowest polygenic load (1^st^ decile) to the highest polygenic load (10^th^ decile). The most significant threshold refers to the P_T_ at which the PGS for SCZ and lithium treatment outcomes were most strongly associated (i.e., the smallest p-value). Using binary logistic and linear regression modeling, we compared BPD patients with lower polygenic load (1^st^ to 9^th^ deciles) for SCZ with patients with the highest polygenic load (10^th^ decile), to quantify the effect of SCZ polygenic load on lithium treatment outcomes. Associations were considered significant at p < 0.05.

The PGS association analyses were adjusted for the covariates age, gender, genotyping platform, and 7 principal components (PCs) calculated in PLINK. The analyses were performed using R for Statistical Computing and PLINK 1.9 for Linux ^37^. Prediction accuracy, the percentage of variance in lithium response accounted by for the PGS at each P_T_, was estimated as the variance explained by the full model including each PGS and covariates minus the variance explained by the model including only covariates.

#### Cross-trait meta-analysis of genome-wide association studies

Biologically, a significantly associated PGS implies that genetic factors influencing the two traits are overlapping. Thus, further analyses were performed to identify genetic polymorphisms that are likely to both increase the susceptibility to SCZ and influence treatment response to lithium in patients with BPD. We performed cross-trait meta-analyses by combining the summary statistics for GWAS on lithium response from the ConLi^+^Gen^22^ and GWAS on SCZ from the PGC^35^. We applied both the O’Brien’s (OB) method and the direct Linear Combination of dependent test statistics (dLC) approach^43,44^, which are implemented in the C^++^ eLX package. Briefly, the OB and dLC approach, combine univariate meta-GWAS data (beta coefficients or Z-scores) for each SNP^43,44^.

The methods follow an inverse-variance meta-analysis approach and directly combine correlated Z-scores (as in meta-analyses) considering the correlation within the univariate test statistics and estimated variances between the traits. The OB method is more powerful when the summary statistics are homogeneous (not very different) and in the similar direction, while dLC is better when the test statistics are either heterogeneous or in opposite directions. Because they often vary based on the sign of the Z-scores, the smallest p-value on either of the two tests could be used to determine statistical significance. Further details are available elsewhere^43,44^.

In this cross-trait meta-analysis, for each SNP we combined GWAS association Z-scores from the SCZ study^35^ with the GWAS association Z-scores for lithium treatment response in the ConLi^+^Gen study^22^ separately for the categorical and continuous outcomes. Each analysis generates two test statistics and associated p-values, one for the OB method and one for the dLC method. Statistical significance of the cross-trait association was determined based on the smaller of the two p-values. The results were considered significant if (1) the p-value for the cross-trait meta-analysis reached genome-wide significance (p< 5×10^−8^), and; (2) the univariate meta-GWAS effects were at least nominally significant for both SCZ and lithium response (p< 0.01). For each cross-trait meta-analysis, only one independent lead SNP per locus was reported. Nearby SNPs in LD (r^2^>0.1) with the lead SNP were considered dependent and belonging to the same locus.

#### Ingenuity^®^ Pathway Analysis (IPA^®^)

To characterize the potential biological significance of the SNPs discovered from the cross-trait meta-analyses, we performed analyses using QIAGEN's Ingenuity^®^ Pathway Analysis (IPA^®^, QIAGEN Redwood City, CA, USA, www.qiagen.com/ingenuity).

To prepare the input genes for IPA, we followed a three-step bioinformatics approach: *Step 1:* We defined tagSNPs that are in high linkage disequilibrium (LD: r^2^>0.5) and within a + 500-kb region with the meta-GWAS significant SNPs (gSNPs) using the genetic catalog of the 1000 Genomes project phase 3, October 2014 release^45^. *Step 2:* The gSNPs and tagSNPs from step 1 were mapped to the genes in which they are located. This generated a list of hosting genes (hGenes). *Step 3:* We performed an expression quantitative trait loci (eQTL) lookup in three databases, searching for any nearby genes (eGenes) whose expression was associated with each of the gSNPs and tagSNPs from step-1. These databases contained the results of eQTL-mapping studies from blood and/or brain tissues: 1) Westra et al^46^ at FDR<0.05 http://genenetwork.nl/bloodeqtlbrowser/, 2) Almanac (Braineac)^47^ at p<1×10^−5^ http://www.braineac.org/, and 3) Genotype-Tissue Expression (GTEx) data release V6p (dbGaP Accession phs000424.v6.p1) accessed from the GTEx Portal on February 8, 2017, at https://www.gtexportal.org/home/.

Finally, the combined list of hGenes and eGenes was used as input into the IPA software after removing gene duplicates. IPA compares the proportion of input genes mapping to a biological pathway to the reference genes list in the ingenuity databases. The significance of the overrepresented canonical pathways and functional networks is determined using the right-tailed Fisher’s exact test and later adjusted for multiple testing using the Benjamini-Hochberg (BH) method^48^. Significant results were determined at BH adjusted P-value <0.01.

## RESULTS

### Sample characteristics and lithium treatment response rates

In total, 3,193 patients with BPD who had undergone lithium treatment and had available genotype and clinical data participated in the study. After QC, 2,586 patients remained for analysis, of whom 2,366 were of European ancestry and the rest Asian. The mean (sd) age of all the patients combined was 47.2 (13.9) years and 2,052 (62.7%) were female. In all, 704 (27.2%) had a good response to lithium treatment (ALDA score ≥7). The mean (sd) ALDA score for all participants was 4.9 (3.1) (Table 1).

**Table 1:** The characteristics of patients with BPAD and outcomes with lithium treatment

| **Patient characteristics** | **Categorical outcome<sup>a</sup>**<br>**Good versus poor response** | **Continuous scale<sup>b</sup>**<br>**ALDA score on subscale A** |
| --- | --- | --- |
| **BPAD patients (N)** | 2,586 | 2,244 |
| **Responders, N (%)** | 704 (27.2) | - |
| **Age at interview, mean (s.d)** | 47.2 (13.9) | 47.4 (13.9) |
| **Sex, Women, N (%)** | 1,478 (57.2%) | 1,291(57.5%) |
| **ALDA scale A score, mean (s.d)** | 6.2(3.0) | 6.3 (3.0) |
| **ALDA scale total B mean (s.d)** | 2.5(1.7) | 2.1 (1.2) |
| **ALDA scale total mean (s.d)** | 4.1(3.2) | 4.5(3.1) |

**Legend:** BPAD: Bipolar affective disorder; <sup>a</sup>Total ALDA score $\geq 7$ was defined as good response; <sup>b</sup>Subjects with total B score $>4$ or who had missing data on the total scores on ALDA subscale A or B were excluded.

### Associations of SCZ PGS with lithium treatment response in BPD patients

At the most significantly associated threshold (P_T_ <5×10^−2^), the PGS for SCZ was strongly associated with lithium treatment response in BPD (p=8×10^−5^) for the categorical outcome on the ALDA scale (Figure 1), explaining 0.8% of the variance. For the continuous outcome (total score on the ALDA subscale A), the direction of association was congruent with the finding on the categorical outcome, but was not statistically significant (p>0.05). The association results of the categorical and continuous outcomes at each threshold levels are detailed in Figure 1. In each threshold, a lower polygenic load for SCZ was associated with a favorable lithium treatment response in patients with BPD (Table 2 & Figure 1).

**Figure 1:**
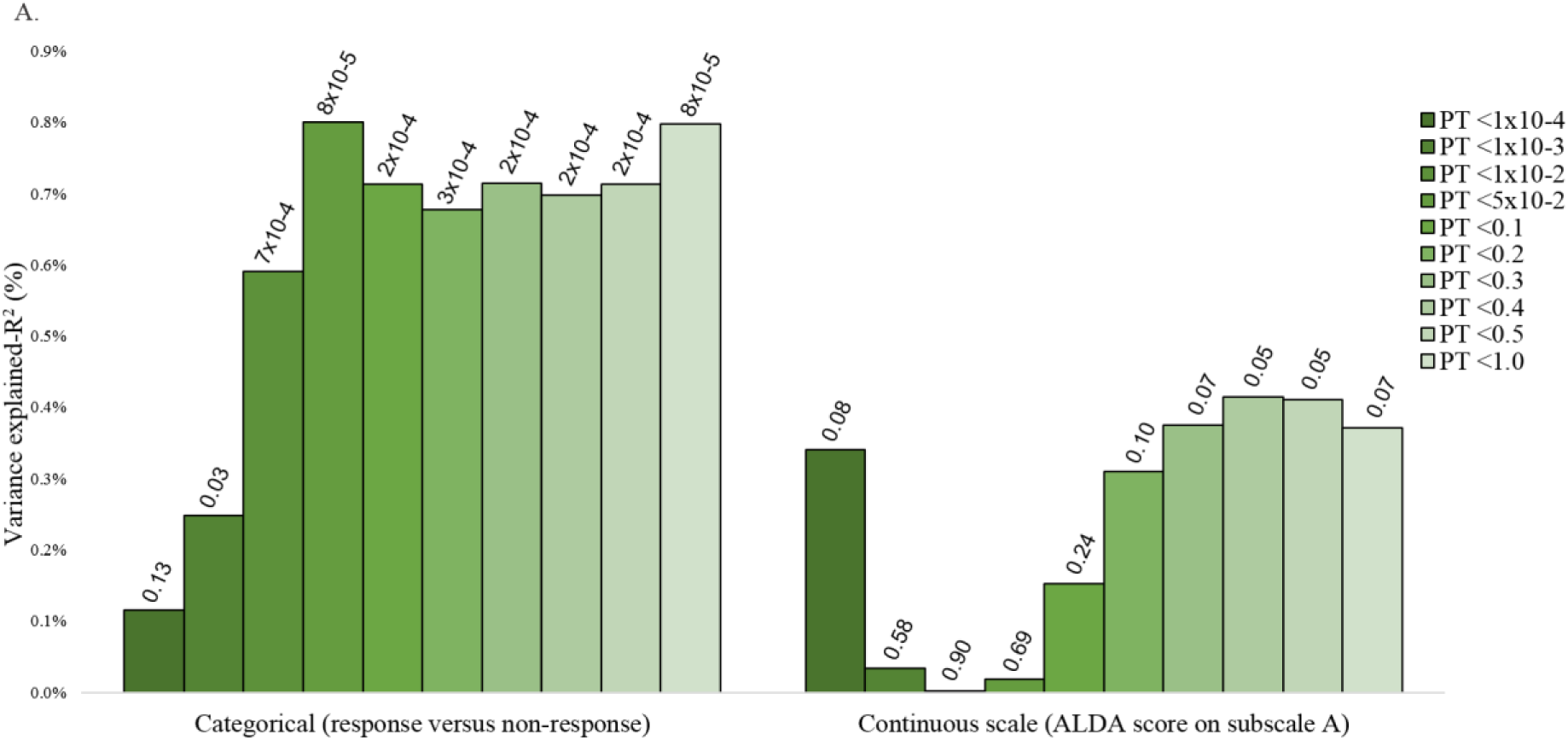
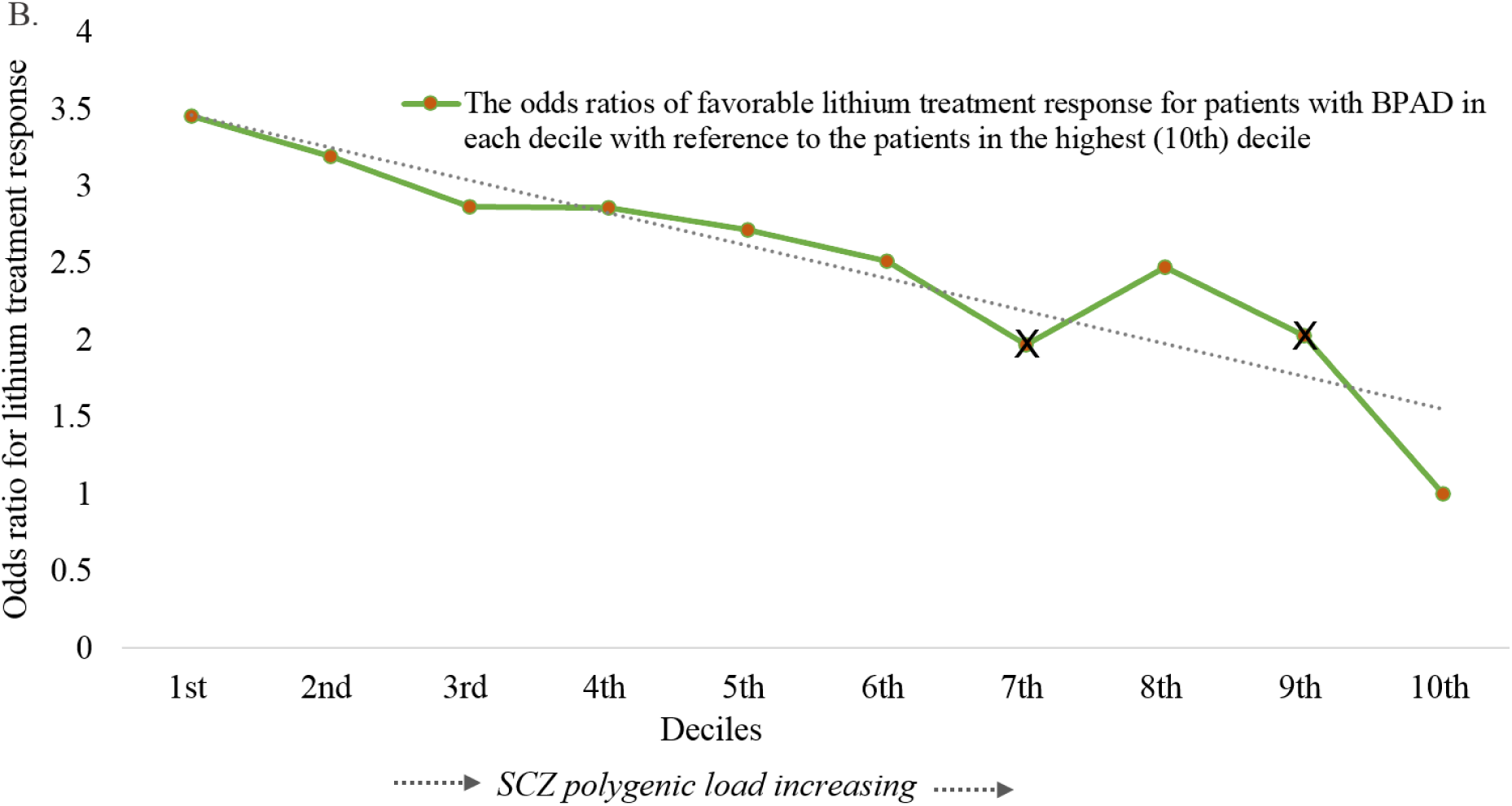
The graph shows (A) the association of polygenic score (PGS) for schizophrenia (SCZ) and lithium treatment response defined as a categorical and continuous scale, at different SCZ GWAS p-value thresholds; and (B) trends in the odds ratios for favorable lithium treatment response for BPD patients in the low SCZ deciles (1^st^ to 9^th^) compared to patients in the highest (10^th^) SCZ PGS decile, estimated at the most significant p-value thresholds (P_T_ <5×10^−2^) (n=2,586). Legend figure 1a: The y-axis (R^2^) refers to the percentage of variance in lithium treatment response accounted for by the PGSs of SCZ at a particular p-value threshold. On the x-axis, plotted from left to right, are the GWAS p-value thresholds used to group single nucleotide polymorphisms (SNPs) for PGSs. On the top of each bar are the p-values of the association between the PGS for SCZ and lithium treatment response. Legend figure 1b: The effect sizes on the y-axis are estimated in odds ratios and on the x-axis are SCZ PGS deciles (1^st^ to 10^th^). **X**-sign on the line plot indicates that the association is not statistically significant at that particular decile.

**Table 2:** The odds ratios of favorable lithium treatment response (categorical outcome) in patients with BPAD, comparing the response status of patients in the low PGS decile for SCZ with patients with the highest polygenic load for SCZ (10^th^ decile).

| SCZ PGS in categories (deciles) | Patients with BPAD (n=2,586) |  |  |
| --- | --- | --- | --- |
|  | <sup>a</sup>R/N | unadjusted OR (95% CI) | <sup>b</sup>Adjusted OR (95% CI) |
| **1<sup>st</sup> lowest score** | 83/175 | 1.97 (1.32-2.96) | 3.46 (1.42-8.41) |
| **2<sup>nd</sup>** | 80/179 | 1.86 (1.24-2.79) | 3.19 (1.32-7.74) |
| **3<sup>rd</sup>** | 78/180 | 1.80 (1.20-2.71) | 2.87 (1.18-6.95) |
| **4<sup>th</sup>** | 76/184 | 1.72 (1.14-2.59) | 2.86 (1.18-6.91) |
| **5<sup>th</sup>** | 76/180 | 1.76 (1.17-2.64) | 2.71 (1.12-6.55) |
| **6<sup>th</sup>** | 67/194 | 1.44 (0.95-2.18) | 2.50 (1.03-6.05) |
| **7<sup>th</sup>** | 58/200 | 1.21 (0.79-1.85) | 1.97 (0.81-4.79) |
| **8<sup>th</sup>** | 75/184 | 1.70 (1.13-2.55) | 2.47 (1.03-5.96) |
| **9<sup>th</sup>** | 61/198 | 1.28 (0.84-1.95) | 2.03 (0.86-4.81) |
| **10<sup>th</sup> highest score** | 50/208 | 1 (reference) | 1 (reference) |

**Legend:** The reference decile refers to the PGS category with the highest polygenic load for schizophrenia (10<sup>th</sup> decile, at $P_T < 5 \times 10^{-2}$ ).
<sup>a</sup> R/N: number of lithium responders versus non-responders; <sup>b</sup> adjusted for age, sex, genotyping platform and 7-principal components. SCZ: schizophrenia, PGS: polygenic score, OR: odds ratio

Table 2 shows the odds ratios (OR) for the association between lithium treatment response in BPD and SCZ PGS in deciles, comparing the response status of patients in the low polygenic load categories (1^st^ to 9^th^ deciles) with patients in the highest polygenic load category for SCZ (in the 10^th^ decile). Results demonstrate that BPD patients who carry a lower polygenic load for SCZ have higher odds of favorable lithium treatment response, compared to patients carrying a high polygenic load. In other words, the OR of favorable treatment response decreased as the genetic load for SCZ increased, ranging from an OR 3.46 [95%CI: 1.42-8.41] at 1^st^ decile to OR 2.03 [95%CI: 0.86–4.81] at the 9^th^ decile, compared to the reference SCZ PGS at the 10^th^ decile. As well, there was a highly significant linear trend in the association between the PGS at deciles and lithium treatment response (Table 1& Figure 1).

### Cross-trait meta-analysis of GWAS for lithium treatment response in BPD, and GWAS for SCZ

Subsequent to the PGS analysis, we performed a SNP-based cross-trait meta-analysis by combining the summary statistics for the GWASs on: 1) SCZ and lithium treatment response in the categorical outcome; and 2) SCZ and lithium treatment response in the continuous outcome - with the aim of identifying individual genetic variants implicated in the genetic susceptibility to SCZ and lithium treatment response. This meta-analysis yielded 15 loci with p-values below the genome-wide significance level (p<5×10^−8^) (Table 3, Figure 2). The top six loci and closest genes were: rs144373461 (p=1.28×10^−17^; *HCG4*), rs66486766 (p=1.38×10^−11^; *ADAMTSL3*), rs7405404 (p=4.62×10^−11^; *ERCC4*), rs142425863 (p=5.13×10^−11^; *HCG4*), rs3919583 (p=4.54×10^−9^; *CCNH*); and rs59724122 (p=5.16×10^−9^; *EPHX2*)

**Table 3:**
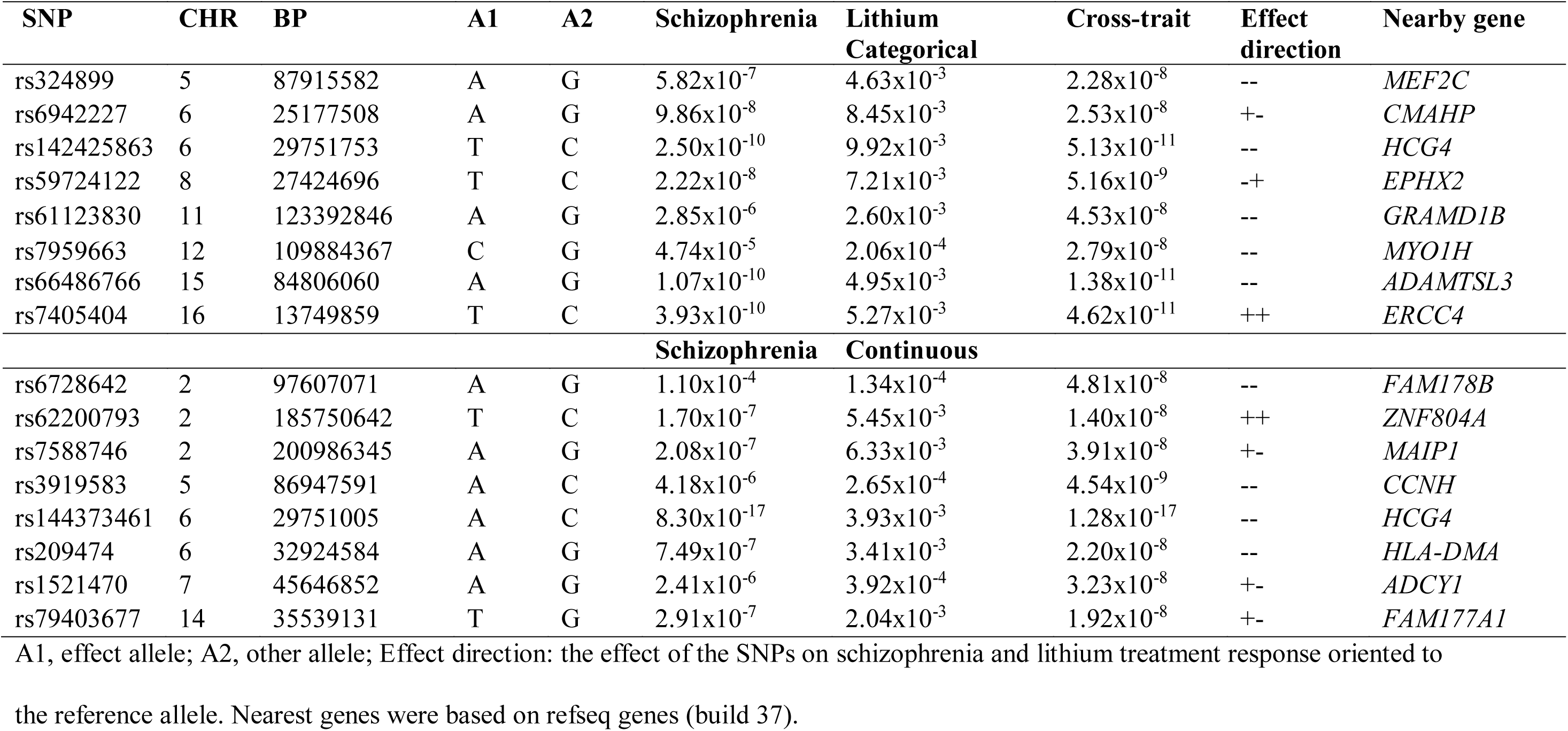
Loci resulting from cross-trait meta-analysis of GWASs for lithium treatment response in BPAD patients and GWAS for SCZ (P-univariateGWAS<1×10-2 and cross-trait P-cross-trait <5×10-8).

**Figure 2:**
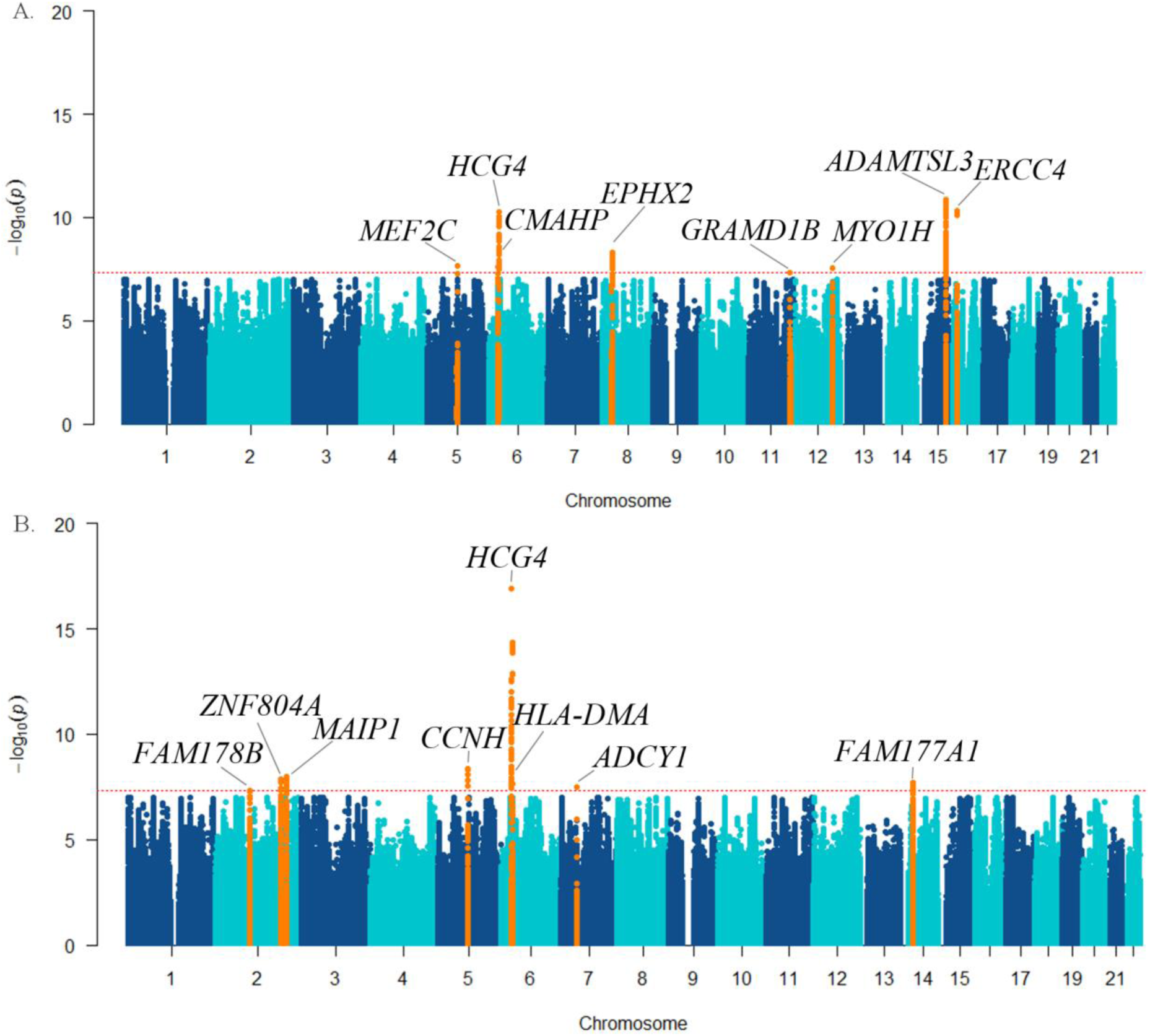
Manhattan plot showing the result of cross-trait meta-analysis of GWASs on SCZ and the GWASs on lithium treatment response in BPD as A) categorical outcome; and B) continuous scale, highlighting the loci that showed genome-wide significance (orange), and the nearest genes (top). Legend figure 2: The —log10 (cross-trait p-value) is plotted against the physical position of each SNP on each chromosome. The threshold for genome-wide significance (cross-trait p-value<5×10^−8^) is indicated by the red dotted horizontal line. To characterize the functional implications of identified SNPs, we undertook IPA pathway analysis using query gene inputs generated from the results of the cross-trait and eQTL analyses. These genes included 33 hGenes hosting the gSNPs and tagSNPs, as well as the eQTL genes identified from the three databases - 27 eGenes from Westra et al, 23 eGenes from Almanac (Braineac) and 31 eGenes in GTEx portal. Table 4 gives the list of 82 unique genes used as input for IPA.

**Table 4:** Combined list of eGenes and hGenes used as an input in the Ingenuity Pathway Analysis (IPA)

| Hosting Genes | eQTL genes (eGenes) lookup in |  |  | All combined Duplicates removed |
| --- | --- | --- | --- | --- |
|  | **Westra** | **BRAINEAC** | **GTEx** |  |
| *HLA-F* | *HLA-G* | *ADAMTSL3* | *HLA-K* | *AC103965.1, ACACB* |
| *HCG4* | *ANKRD36* | *EPHX2* | *HLA-F-AS1* | *ADAMTSL3, ADCY1* |
| *HLA-DMB* | *SRP54* | *HLA-F* | *HLA-H* | *ANKRD36, BAZ1A* |
| *HLA-DMA* | *UBE3B* | *IFITM4P* | *HLA-F* | *BRD2, CASP14* |
| *BRD2* | *KCTD10* | *MMAB* | *ZFP57* | *CHRNA2, CMAHP* |
| *FAM178B* | *ACACB* | *NACAD* | *HLA-V* | *CSPG4P11, EFTUD1P1* |
| *ANKRD36* | *NMB* | *SEC11A* | *HCG4P11* | *EPHX2, FAM177A1* |
| *ZNF804A* | *KIAA0391* | *TRIM26* | *PPP2R3C* | *FAM178B, GABBR1* |
| *TMEM161B* | *PPP1R11* | *TRIM35* | *CSPG4P11* | *GOLGA6L4, GOLGA6L5P* |
| *MEF2C* | *HLA-F* | *UBE3B* | *ZSCAN2* | *GPNMB, HCG4* |
| *ADCY1* | *ZNRD1* | *WDR73* | *HLA-W* | *HCG4B, HCG4P11, HCG4P5* |
| *MYO1H* | *MMAB* | *ZNF592* | *IFITM4P* | *HFE, HLA-A, HLA-DMA* |
| *KCTD10* | *EPHX2* | *HCG4B* | *HLA-A* | *HLA-DMB, HLA-DOB, HLA-DPB1* |
| *UBE3B* | *TAP2* | *ZNRD1ASP* | *HLA-U* | *HLA-F, HLA-F-AS1, HLA-G* |
| *MMAB* | *HLA-DPB1* | *SCAND2P* | *HLA-J* | *HLA-H, HLA-J, HLA-K, HLA-T* |
| *MVK* | *HLA-DMB* | *KIAA1920* | *MICF* | *HLA-U, HLA-V, HLA-W, IFITM4P* |
| *IGBP1P1* | *HLA-DMA* | *LOC100128364* | *MVK* | *IGBP1P1, KCTD10, KIAA0391* |
| *SRP54* | *HLA-DOB* | *LOC285830* | *EFTUD1P1* | *KIAA1920, LMAN2L* |
| *FAM177A1* | *LMAN2L* | *LOC440297* | *GOLGA6L4* | *LOC100128364, LOC285830* |
| *PPP2R3C* | *GABBR1* | *LOC440300* | *CHRNA2* | *LOC440297, LOC440300* |
| *KIAA0391* | *PSMB9* | *LOC642288* | *IGBP1P1* | *LOC642288, LOC727858* |
| *PSMA6* | *BRD2* | *LOC727858* | *BAZ1A* | *LOC728121, MEF2C, MICD* |
| *ADAMTSL3* | *MEF2C* | *LOC728121* | *MICE* | *MICE, MIC, MMAB, MVK* |
| *GOLGA6L5P* | *PPP2R3C* |  | *HCG4P5* | *MYO1H, NACAD, NMB* |
| *UBE2Q2P1* | *CMAHP* |  | *MICD* | *PPP1R11, PPP2R3C, PSMA6* |
| *ZSCAN2* | *GPNMB* |  | *HLA-G* | *PSMB9, SCAND2P, SEC11A* |
| *SCAND2P* | *SEC14L3* |  | *HLA-T* | *SEC14L3, SRP54, TAP2* |
| *WDR73* |  |  | *GOLGA6L5P* | *TMEM161B, TRIM26, TRIM35* |
| *NMB* |  |  | *HFE* | *UBE2Q2P1, UBE3B* |
| *SEC11A* |  |  | *CASP14* | *WDR73, ZFP57, ZNF592* |
| *ZNF592* |  |  | *AC103965.1* | *ZNF804A, ZNRD1* |
| *GPNMB* |  |  |  | *ZNRD1ASP, ZSCAN2* |
| *LOC440300* |  |  |  |  |

We then assessed how these genes are enriched with canonical pathways in the Ingenuity database. The most significantly represented canonical pathways and enriched genes are shown in Table 5. The top 5 IPA^®^ canonical pathways include: *Antigen Presentation Pathway*, *OX40 Signaling Pathway*, *Autoimmune Thyroid Disease Signaling*, *Cdc42 Signaling*, and *B Cell Development* (Table 5). These pathways were predominantly identified on the basis of several HLA genes —*HLA-A*, *HLA-DMA*, *HLA-DMB*, *HLA-DOB*, *HLA-DPB1*, *HLA-F*, *HLA-G*, *PSMB9,* and *TAP2*.

**Table 5:**
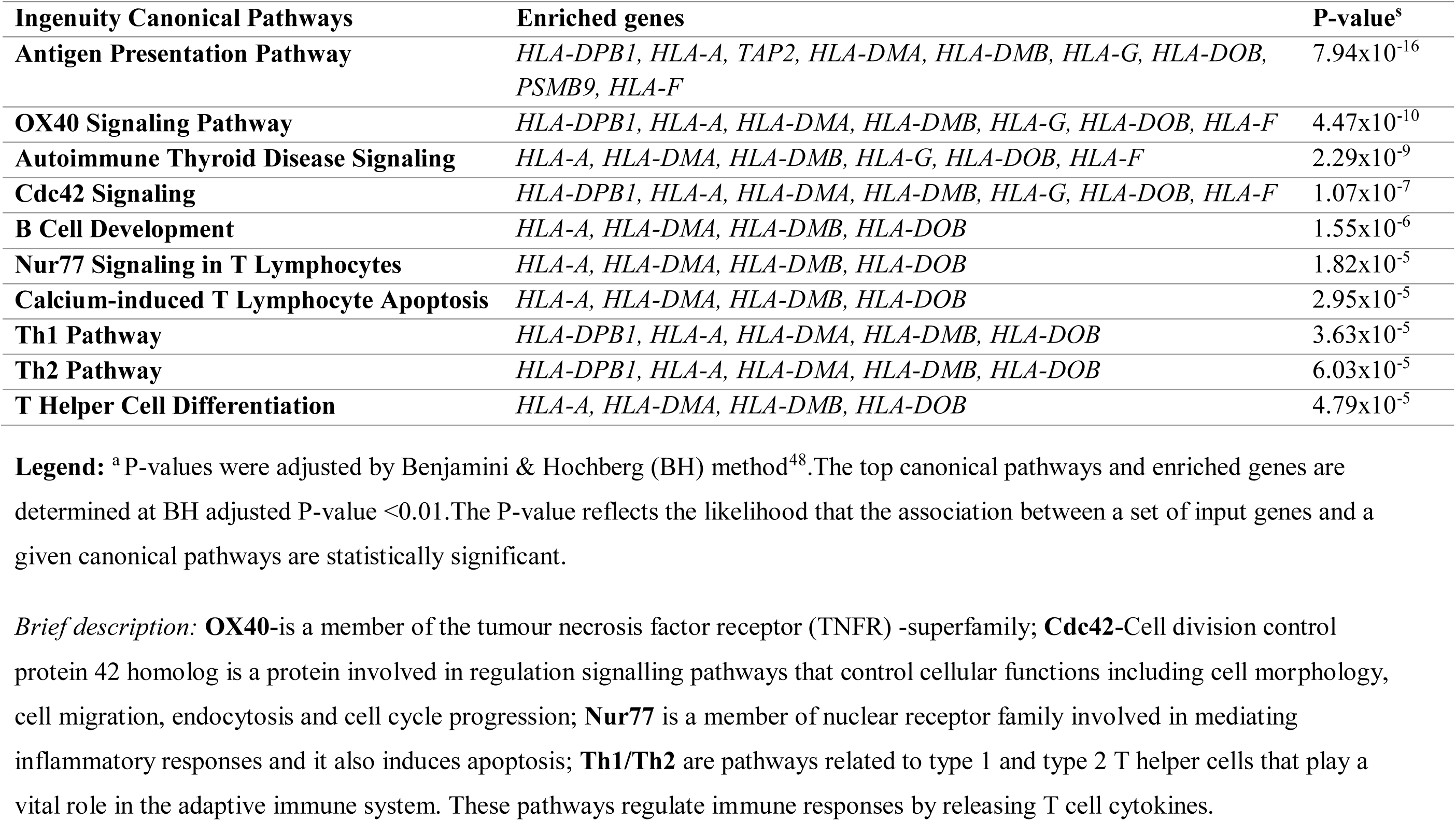
The top canonical signaling pathways enriched for genes identified in the cross-trait meta-analyses

The IPA^®^ network analysis revealed 2 relevant functional networks (Table 6). As it can be seen in Figure 3, the top 2 networks indicate that tumor necrosis factor alpha (TNFα), Interleukin-4 (IL-4), and interferon gamma (IFNγ) might represent important functional molecular nodes in the interaction between lithium response and SCZ.

**Figure 3:**
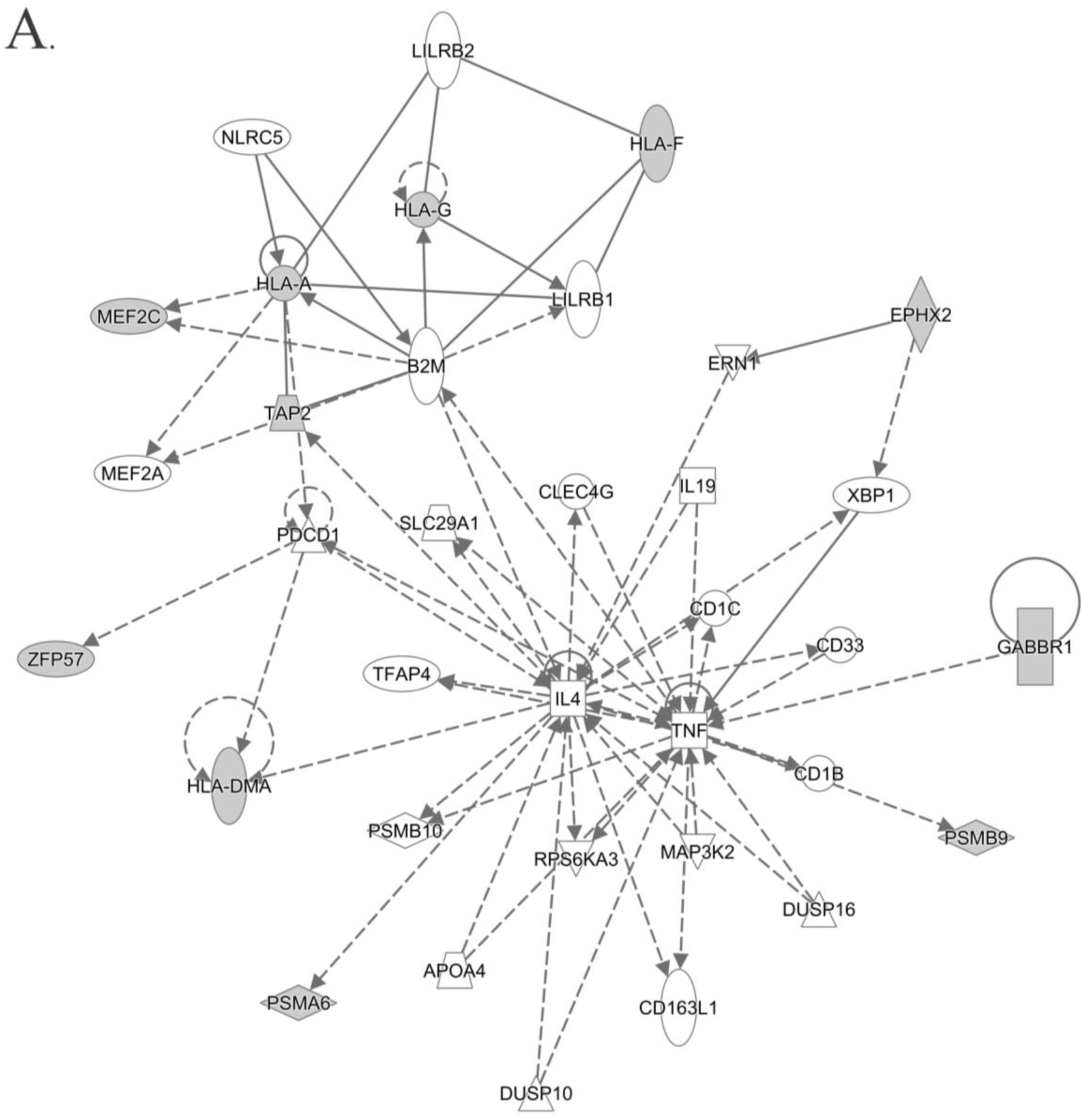
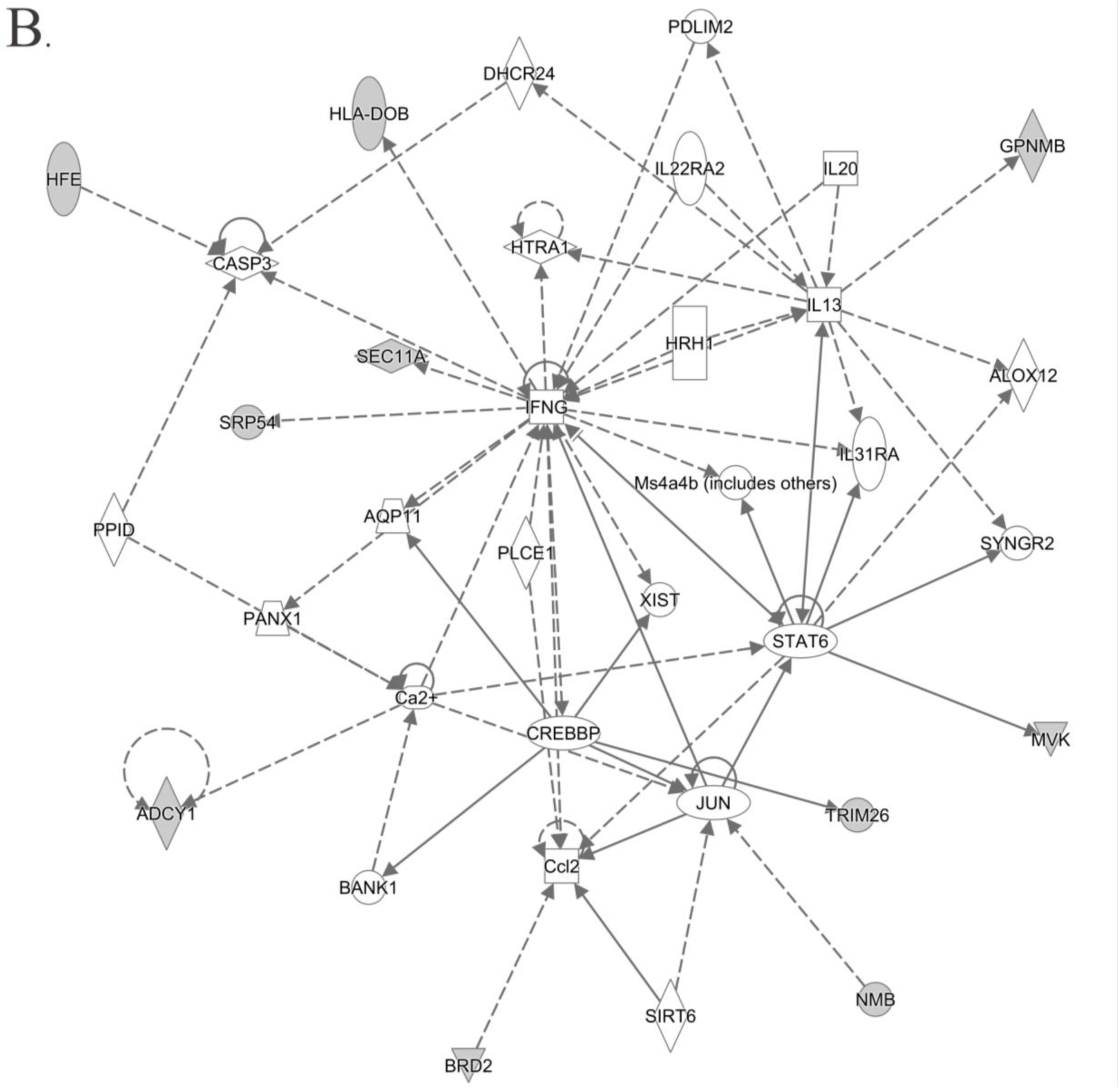
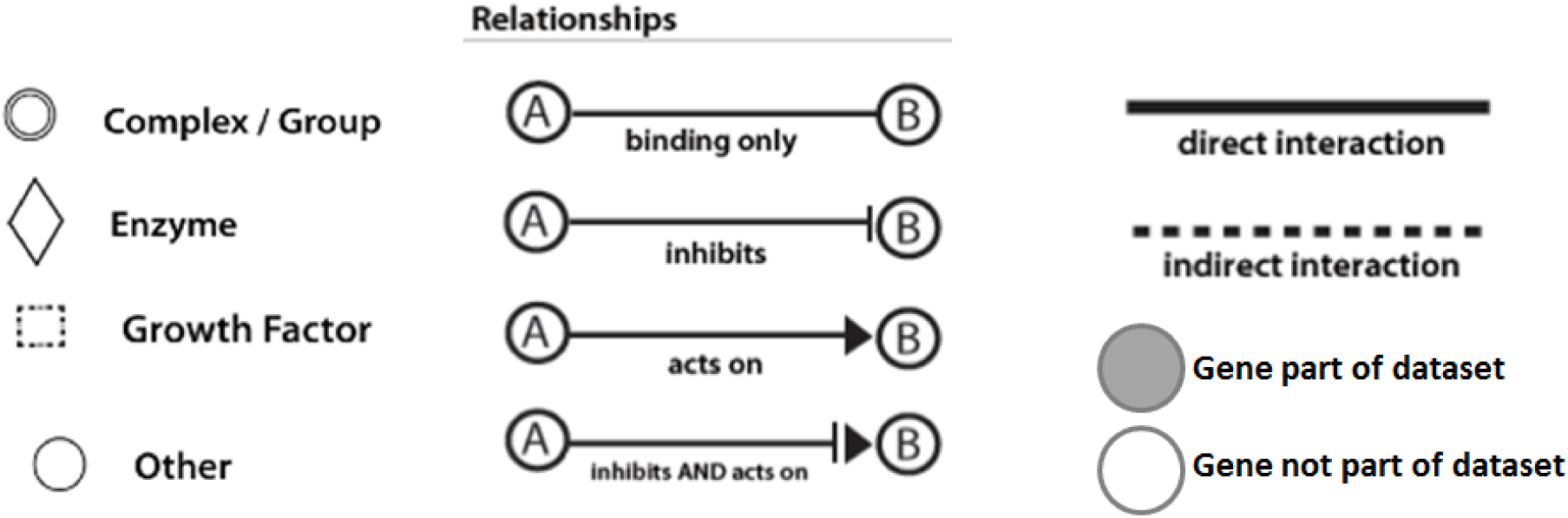
Indicates the top networks of molecules in IPA, in which TNFα, IL-4 and IFNG represent the main functional nodes mediating the genetic interaction between lithium response and SCZ; shown as network for a) group 1 and b) group 2 molecules in Table 6. Legend Figure 3: IPA generates the network using a proprietary algorithm, and included genes that could contribute to the network, even if they were not contained in the original dataset.

**Table 6:**
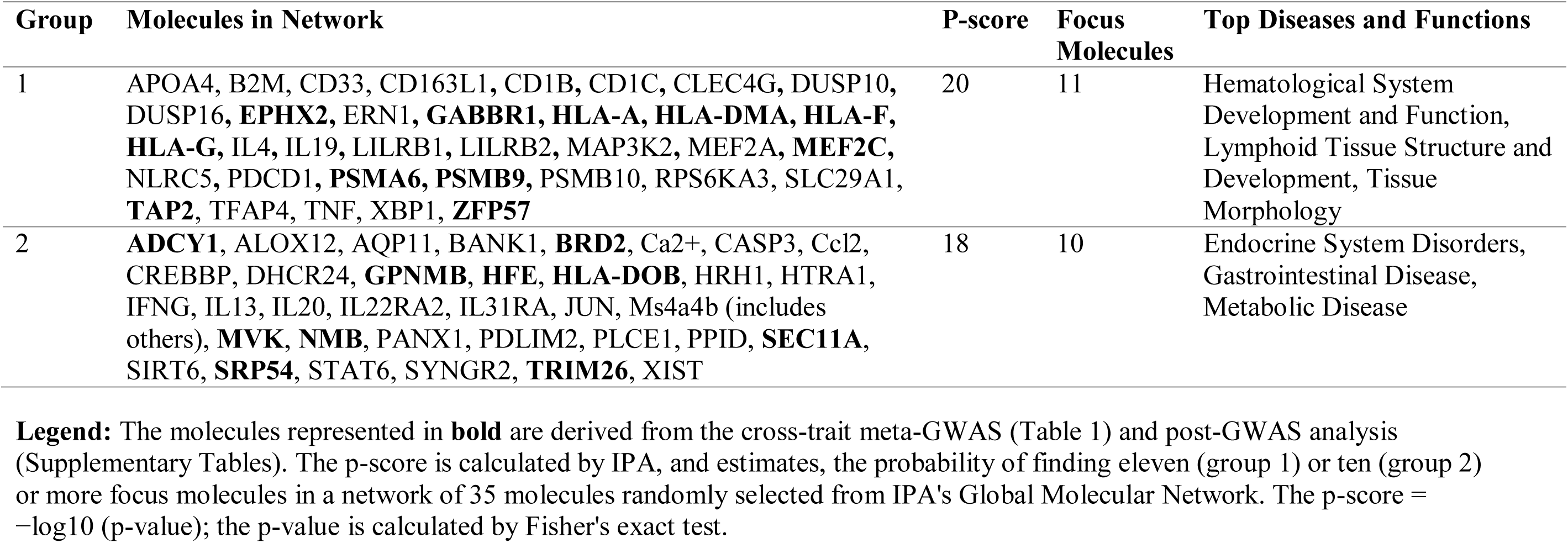
Top IPA protein networks, molecules in network and top diseases functionally related to the network

| Group | Molecules in Network | P-score | Focus Molecules | Top Diseases and Functions |
| --- | --- | --- | --- | --- |
| 1 | APOA4, B2M, CD33, CD163L1, CD1B, CD1C, CLEC4G, DUSP10, DUSP16, **EPHX2**, ERN1, **GABBR1**, **HLA-A**, **HLA-DMA**, **HLA-F**, **HLA-G**, IL4, IL19, LILRB1, LILRB2, MAP3K2, MEF2A, **MEF2C**, NLRC5, PDCD1, **PSMA6**, **PSMB9**, PSMB10, RPS6KA3, SLC29A1, **TAP2**, TFAP4, TNF, XBP1, **ZFP57** | 20 | 11 | Hematological System Development and Function, Lymphoid Tissue Structure and Development, Tissue Morphology |
| 2 | **ADCY1**, ALOX12, AQP11, BANK1, **BRD2**, Ca2+, CASP3, Ccl2, CREBBP, DHCR24, **GPNMB**, **HFE**, **HLA-DOB**, HRH1, HTRA1, IFNG, IL13, IL20, IL22RA2, IL31RA, JUN, Ms4a4b (includes others), **MVK**, **NMB**, PANX1, PDLIM2, PLCE1, PPID, **SEC11A**, SIRT6, **SRP54**, STAT6, SYNGR2, **TRIM26**, XIST | 18 | 10 | Endocrine System Disorders, Gastrointestinal Disease, Metabolic Disease |

## DISCUSSION

The present study reports two main findings: first, using PGS methodology, we demonstrate that there is an inverse association between genetic loading for SCZ risk variants and long-term therapeutic response to lithium in patients with BPD on the categorical outcome of the ALDA scale. Second, we show in cross-trait meta-GWAS and pathway analyses that genetic variants in the HLA region, the antigen presentation pathway and inflammatory cytokines such as TNF-α, IL-4 and IFNγ could have a biological role in lithium treatment response in BPD.

These findings are consistent with previous clinical and epidemiological studies of lithium response. Lithium is not an effective medication for people suffering from SCZ spectrum disorders^49,31^. Moreover, lithium may be deleterious for patients with SCZ because of their greater liability to developing lithium-induced neurotoxicity even at modest doses and blood levels^49,50^. The severity of psychotic symptoms present in bipolar patients was found inversely associated with lithium treatment response^51^. Similarly, slow resolution of psychosis in response to lithium treatment during acute manic episodes has been shown to predict poorer overall response to the drug^52^. Amongst patients with BPD, those with a family history of SCZ show poorer response to lithium compared to those with a family history of BPD^53^. Our findings may provide insight into the genetic architecture underlying these clinical observations.

In the SCZ to lithium response cross-trait GWAS meta-analyses, 15 genetic loci located within protein-coding, genes that appear to have overlapping effects on SCZ risk and response to lithium treatment in BPD were identified. Only one of these genes, type 1 adenylyl cyclase *(ADCY1)*, had previously been directly implicated in genetic studies of both SCZ^54^ and lithium treatment response^26^. It has been shown that *ADCY1* directs neuronal signaling through activation of the extracellular signal-regulated kinases 1 and 2 (ERK1/2) and phosphoinositide 3-kinase (PI3K) pathways^55^. Lithium, in turn, has also been shown to engage the ERK 1/2 pathway and the PI3K pathway, possibly through complex interactions with GSK-3^56,57^. It is possible that the polymorphisms in the *ADCY1* gene implicated in our study result in altered ERK1/2 and PI3K activation states, thereby interfering with potentially therapeutic lithium effects through these pathways.

Both the most significant finding of the cross-trait GWAS *(HCG4* gene on chromosome 6) and the SNPs from the post-GWAS functional analyses point out to the HLA system in modulating lithium response. Differences in cellular HLA surface protein composition between BPD patients who respond well to lithium and non-responders were first reported over 30 years ago in several studies. These reports noted that leukocyte HLA-A3 antigen reactivity was reported to be associated with poor lithium response, whereas the absence of HLA-A3 predicted a favorable response^58–60^. At the same time, *in vitro* experiments suggested that lithium binds to HLA antigens on cultured human leukocytes^61^. A subsequent *in vivo* study in BPD patients demonstrated that exposure to lithium for about 2 months promoted substantial alterations in the composition of leukocyte HLA proteins^62^.

The genetic association between SCZ and the HLA region on chromosome 6 is the most robust finding of SCZ GWAS to date^35,63–66^. A functional follow-up analysis demonstrated that HLA SCZ risk variants result in altered expression of Complement Component 4a (C4a) and 4b (C4b) proteins, impacting negatively on neuronal synaptic pruning and thereby resembling neuropathological findings in SCZ^67^. Further, reduced C4a in transgenic mice resulted in greatly decreased neuronal complement component 3 (C3) expression^67^. Interestingly, a recent study demonstrated that lithium exposure of human monocytes and mouse microglia *in vitro* resulted in increased expression of C3, which in turn was driven by the inhibition of glycogen synthase kinase-3 (GSK-3)^68^. Inhibition of GSK-3 is to date the most comprehensively documented molecular effect of lithium in neurons, glia, and peripheral immune cells^69,70^. Taken together, these studies and our findings raise the possibility that lithium's GSK-3-mediated activation of the complement system, via enhanced C3 expression^68^, is suppressed in people with a high genetic loading for SCZ due to functional disturbances of the complement cascade resulting from the SCZ-HLA-C4 association. In this context, it is also compelling that IPA^®^ identified Antigen Presentation as the top canonical pathway characterizing the findings of our cross-trait meta-GWAS analysis. Cellular antigen presentation is mediated by HLA proteins and is closely linked to the functions of the complement system as described above.

Further, functional network analysis of our meta-GWAS findings implicated TNFα, IL-4 and IFN-γ as central functional nodes, suggesting that the negative interaction between lithium response and genetic predisposition for SCZ could be mediated by mechanisms implicating these pro-inflammatory cytokines. Previous studies have reported modulatory effects of lithium treatment on these cytokines in BPD. For example, a study of euthymic patients with BPD reported that TNFα and IL-4 were selectively increased in patients on lithium monotherapy relative to untreated patients and healthy controls^71^. Similarly, lithium treatment in BPD patients with a rapid cycling pattern was associated with increased TNFα levels^72^. In a large clinical sample, peripheral TNFα activity was increased in people with SCZ and BPD, and there was evidence that lithium treatment further increased serum levels in those with BPD^73^. In contrast, *in vitro* experiments have shown that lithium decreases IFN-γ levels in human blood cultures^74,75^, while attenuating the differentiation of naïve CD4^+^ T cells into T1 Helper (Th1) cells by IFN-γ following immune challenge^76^. These effects on inflammatory cytokines are, at least in part, driven by GSK-3 inhibition^77,76,78^. Intriguingly, one study using the ALDA scale reported elevated TNFα levels in patients with poor long-term response to lithium, compared to good responders^79^. In all, these findings underscore the possibility that mechanisms involving pro-inflammatory cytokines might play an important role in mediating therapeutic effects of lithium in patients with BPD^80^. The disturbances of these mechanisms through genetic variants involved in the pathogenesis of SCZ might also perturb lithium's clinical effectiveness. A growing body of evidence describing aberrant inflammatory processes in patients with first episode psychosis^81^ and SCZ^82^ supports this idea.

This study has four limitations that are outlined in the Supplementary Materials.

### Limitations of the study

Our study has a number of limitations. First, the polygenic load for SCZ accounted for only a modest percentage (~1%) of the observed variation in lithium treatment response in patients with BPD. While this is in line with previous reports on the effects of PGSs on complex clinical phenotypes such as SCZ and BPD^65^, the significance of this finding at clinical- and population-levels needs to be further explored. Encouragingly, previous studies indicate that PGS approaches can assist in characterizing relevant clinical phenotypes. For example, in SCZ, a high polygenic SCZ score has been reported as a measure of disease chronicity^83^, and is associated with failure to respond to treatment^84^. Second, lithium response in our study was assessed using the ALDA scale, which is a retrospective measure.

In order to substantiate our findings further, prospective studies are required that can measure clinical responses to lithium prospectively. Third, while our strategy for exploring the biological context of our genetic findings can point towards avenues for future research, it is not designed to provide definitive mechanistic answers. Hypothesis-driven experiments are required to follow up on these leads. Fourth, the Ingenuity Pathway Analysis revealed that the enriched pathways were mainly driven by two independent loci (rs209474 and rs144373461/rs142425863). As an example, the top associated "Antigen Presentation Pathway" contains a total of 9 genes of which 6 are implicated by the SNP rs209474 (*HLA-DPB1, TAP2, HLA-DMA, HLA-DMB, HLA-DOB, and PSMB9*, all genes located at chr6:32,768,557-33,059,376, hg19) and the other 3 genes (*HLA-A, HLA-F, HLA-G*, all located at chr6:29,683,619-29,917,908) by the SNPs rs144373461 and rs142425863 which have a chromosomal distance of only 748 bp. This could be due to the high LD structure in the HLA region and also be related to the parameters used to define LD to extract tagSNPs to the meta-GWAS significant SNPs (LD: r^2^>0.5 and within a + 500-kb region). The same commonly used parameters were used for all significant findings without a priori stratification according to a chromosomal region.

In conclusion, we demonstrated for the first time that lower SCZ loading is strongly associated with better lithium response in patients with BPD. Follow-up functional analyses point to genes that code for the immune system, including the HLA complex and inflammatory cytokines. For future clinical translation, a high genetic loading for SCZ risk variants could be used in conjunction with clinical parameters to predict the likelihood of non-response to lithium treatment in BPD.

## Conflict of interest

All authors declare they have no competing interests

## Acknowledgment

The authors are grateful to all patients who participated in the study and we appreciate the contributions of clinicians, scientists, research assistants and study staff who helped in the patient recruitment, data collection and sample preparation of the studies. We are also indebted to the members of the ConLi^+^Gen Scientific Advisory Board (http://www.conligen.org/) for critical input over the course of the project.

The analysis of this study was carried out using the high-performance computational capabilities of the University of Adelaide, Phoenix supercomputer https://www.adelaide.edu.au/phoenix/ and Lisa Computer Cluster within the Dutch national e-infrastructure (www.surfsara.nl).

## Funding

The primary sources of funding were the Deutsche Forschungsgemeinschaft (DFG; grant no.RI 908/7-1; grant FOR2107, RI 908/11-1 to Marcella Rietschel, NO 246/10-1 to Markus M. Nöthen) and the Intramural Research Program of the National Institute of Mental Health (ZIA-MH00284311; ClinicalTrials.gov identifier: NCT00001174).

The genotyping was in part funded by the German Federal Ministry of Education and Research (BMBF) through the Integrated Network IntegraMent (Integrated Understanding of Causes and Mechanisms in Mental Disorders), under the auspices of the e:Med Programme (grants awarded to Thomas G. Schulze, Marcella Rietschel, and Markus M. Nöthen).

Some data and biomaterials were collected as part of eleven projects (Study 40) that participated in the National Institute of Mental Health (NIMH) Bipolar Disorder Genetics Initiative. From 2003-2007, the Principal Investigators and Co-Investigators were: Indiana University, Indianapolis, IN, R01 MH59545, John Nurnberger, M.D., Ph.D., Marvin J. Miller, M.D., Elizabeth S. Bowman, M.D., N. Leela Rau, M.D., P.Ryan Moe, M.D., Nalini Samavedy, M.D., Rif El-Mallakh, M.D. (at University of Louisville), Husseini Manji, M.D.(at Johnson and Johnson), Debra A.Glitz, M.D.(at Wayne State University), Eric T.Meyer, Ph.D., M.S.(at Oxford University, UK), Carrie Smiley, R.N., Tatiana Foroud, Ph.D., Leah Flury, M.S., Danielle M.Dick, Ph.D (at Virginia Commonwealth University), Howard Edenberg, Ph.D.; Washington University, St. Louis, MO, R01 MH059534, John Rice, Ph.D, Theodore Reich, M.D., Allison Goate, Ph.D., Laura Bierut, M.D.K02 DA21237; Johns Hopkins University, Baltimore, M.D., R01 MH59533, Melvin McInnis, M.D., J.Raymond DePaulo, Jr., M.D., Dean F. MacKinnon, M.D., Francis M. Mondimore, M.D., James B. Potash, M.D., Peter P. Zandi, Ph.D, Dimitrios Avramopoulos, and Jennifer Payne; University of Pennsylvania, PA, R01 MH59553, Wade Berrettini, M.D., Ph.D.; University of California at San Francisco, CA, R01 MH60068, William Byerley, M.D., and Sophia Vinogradov, M.D.; University of Iowa, IA, R01 MH059548, William Coryell, M.D., and Raymond Crowe, M.D.; University of Chicago, IL, R01 MH59535, Elliot Gershon, M.D., Judith Badner, Ph.D., Francis McMahon, M.D., Chunyu Liu, Ph.D., Alan Sanders, M.D., Maria Caserta, Steven Dinwiddie, M.D., Tu Nguyen, Donna Harakal; University of California at San Diego, CA, R01 MH59567, John Kelsoe, M.D., Rebecca McKinney, B.A.; Rush University, IL, R01 MH059556, William Scheftner, M.D., Howard M. Kravitz, D.O., M.P.H., Diana Marta, B.S., Annette Vaughn-Brown, M.S.N., R.N., and Laurie Bederow, M.A.; NIMH Intramural Research Program, Bethesda, MD, 1Z01MH002810-01, Francis J. McMahon, M.D., Layla Kassem, Psy.D., Sevilla Detera-Wadleigh, Ph.D, Lisa Austin, Ph.D, Dennis L. Murphy, M.D.; Howard University, William B. Lawson, M.D., Ph.D., Evarista Nwulia, M.D., and Maria Hipolito, M.D. This work was supported by the NIH grants P50CA89392 from the National Cancer Institute and 5K02DA021237 from the National Institute of Drug Abuse.

The Canadian part of the study was supported by the Canadian Institutes of Health Research to MA grant #64410 to MA. Collection and phenotyping of the Australian UNSW sample, by Philip B. Mitchell, Peter R. Schofield, Janice M. Fullerton and Adam Wright, was funded by an Australian NHMRC Program Grant (No.1037196). The collection of the Barcelona sample was supported by the Centro de Investigación en Red de Salud Mental (CIBERSAM), IDIBAPS, and the CERCA Programme / Generalitat de Catalunya (grant numbers PI080247, PI1200906, PI12/00018, 2014SGR1636, and 2014SGR398). The Swedish Research Council, the Stockholm County Council, Karolinska Institutet and the Söderström-Königska Foundation supported this research through a grant by Lena Backlund. The collection of the Geneva sample was supported by the Swiss National Foundation (grants Synapsy 51NF40-158776 and 32003B-125469). The collection of the Romanian sample was supported by U.E.F.I.S.C.D.I., Romania, grant awarded to Maria Grigoroiu-Serbanescu.

## Web resources

The URLs for data presented herein are as follows:
PGC-Psychiatric Genomics Consortium: schizophrenia, GWAS data, http://www.med.unc.edu/pgc/downloads Blood eQTL browser: http://genenetwork.nl/bloodeqtlbrowser The Brain eQTL Almanac (Braineac): http://www.braineac.org/ The Genotype-Tissue Expression (GTEx): http://www.gtexportal.org/home/.

## Tools

OB and dLC methods in eLX package: https://sites.google.com/site/multivariateyihsianghsu/.

